# A decision matrix to better identify repeatable physiological variation within individuals

**DOI:** 10.1101/2025.07.14.664826

**Authors:** Yangfan Zhang, Chris M. Wood, Colin J. Brauner, Anthony P. Farrell

## Abstract

The performance of an individual has remained at the heart of evolutionary biology since the time of Darwin. Physiologists are equally drawn to the implications of individual variation for health and sporting endeavours, and specifically, whether or not a physiological trait is repeatable within an individual. Experimental biologists are especially interested in temporally stable physiological traits that are relevant to an individual’s lifetime fitness for natural selection to act upon. Experimental noise, however, confounds the measurement of such repeatability, even though validated protocols exist for measuring many meaningful physiological performance traits. Missing is a decision matrix that helps distinguish individual variation from experimental noise. We propose a precision-&-repeatability assessment matrix (PRAM) that integrates established assessments of individual variability and repeatability. This matrix places metrics that are more repeatable and precise in the quadrant closest to the origins of Cartesian coordinates; those farthest away are less acceptable in terms of both repeatability and precision. As a case study, PRAM is applied to whole-organism aerobic and non-aerobic metabolic performance metrics from fish that were measured with the same protocols. The analysis illustrates that aerobic metabolic metrics can be more repeatable and precise than non-aerobic ones. Consequently, PRAM helps physiologists to better understand whether the observed variability is due to non-repeatable metrics or true individual variation.

## Glossary

Absolute aerobic scope (AAS): the numerical difference between maximum and standard metabolic rate, defining the aerobic capacity of animals for their activities [1].

Aerobic metabolism: metabolic pathways supported by oxidative phosphorylation under a steady-state condition.

Aerobic metric: a metric measured with a standardized test at the organismic level that estimates either the rate, performance or capacity of aerobic metabolism.

Accumulated oxygen deficit (AOD): Accumulated oxygen deficit is the anaerobic capacity of an animal in oxygen equivalents to be comparable with absolute aerobic scope [2]. It can be measured as the cumulative oxygen deficit when animals are in a severely low oxygen level below critical oxygen saturation and before the animals lose their upright equilibrium.

Anaerobic metabolism: metabolic pathways temporarily supported by substrate-level phosphorylation. Also referred to as non-aerobic at the whole-animal level.

Non-aerobic metric: a metric measured with a standardized test at the organismic level that reflects the performance or capacity of anaerobic pathways. The metric is measured at the organismic level with a standardized test that reflects the performance and capacity based on high-energy phosphate stores, oxygen stores in tissues and substrate-level phosphorylation metabolic pathways.

Bland-Altman analysis: an established statistical method that quantifies measurement variability when a metric is remeasured on the same individual [3].

Critical oxygen saturation: minimum O_2_ saturation (critical O_2_ saturation, O_2crit_) needed to sustain standard metabolic rate [4] [5].

Ectotherm: an animal whose body temperature conforms to ambient temperature.

Experimental noise: experimental noise that is attributable to operational, instrumental, methodological, and/or analytical variability. Experimental noise is reduced through better controls (either methodologically or statistically), better testing equipment and better training of personnel who make the measurements.

Excess post-exercise oxygen consumption (EPOC): O_2_ requirement to restore tissue O_2_ stores and high-energy phosphate stores, and to metabolize the end products of substrate-level phosphorylation after exhaustive exercise [6]. EPOC is another form of accumulated oxygen deficit.

Factorial aerobic scope (FAS): Factorial aerobic scope measures the aerobic capacity as the ratio to the minimum maintenance oxygen demands (maximum O_2_ uptake / standard metabolic rate) [7].

Factorial scope for oxygen deficit (FSOD): the ratio between critical oxygen saturation and incipient lethal oxygen saturation [8].

Incipient lethal oxygen saturation (ILOS): oxygen level when the fish loses its upright equilibrium, an assessment of hypoxia tolerance [9].

Individual repeatability: The degree to which a trait in an individual is statistically similar each time it is remeasured. While individual repeatability can be statistically tested, repeatability is typically considered within a specific and defined timescale.

Individual variability: The degree to which a trait numerically varies each time it is remeasured in an individual; The converse of individual repeatability.

Maximum oxygen consumption rate (*Ṁ*O_2max_): the maximum attainable rate of oxygen uptake [6] [10].

Physiological metric: a physiological measurement made at one or multiple levels of a biological system.

Precision: the variability of a repeated measurement; higher numerical variability reflects less precision. Therefore, identifying or improving the precision of measurement improves our confidence in assessing the repeatability of the trait by reducing the risk of Type-II statistical errors so that true biological variation can be detected [11].

Routine metabolic rate (RMR): metabolic rate when animals are in a state of routine activities, which is very much situation-specific [12].

Scope for oxygen deficit (SOD): the numerical difference between critical oxygen saturation and incipient lethal oxygen saturation [8].

Stability: A condition of a metric or trait measured in an individual that exhibits high repeatability. A less variable trait has greater stability. Stability (*see* below) is inferred from a statistical test of a metric’s repeatability within an individual.

Standard metabolic rate: the minimum maintenance metabolic rate in the postabsorptive state.

Time instantaneous *Ṁ*O_2_ is above 50% absolute aerobic scope (T_>50%AAS_): The amount of time an animal spends above 50% of its absolute aerobic scope during measurement of standard metabolic rate [13].

Trait: a well-defined biological metric that has proven mechanistic underpinnings.

Variation: describes the differences in values for a given metric among individuals [14] [15]. Variation is a theoretical concept of frequency distribution pertaining to true biological differences seen among individuals within a population for a metric.

Variability: Variability combines biological variation among a group of individuals and the influence of experimental noise on the values for a given metric, *i.e.*, the full tendency of individual differences in a metric [14] [15] [16].

* *The abbreviations listed in the glossary will be directly referred to in the main text*.

## Conceptualizing the challenge

Individual variation in physiological performance traits is fundamental for evolution through natural selection [17] [18]. Thus, the performance of individuals ultimately influences the chance of success of a species, a strain, or a population, just as individual human athletes succeed in sports competitions. Consequently, the past century of science has emphasized the importance of individual variation in evolutionary processes [19], and individual-based models have become central in ecology for many decades [20] [21].

Likewise, performance studies in elite athletes have revealed why ‘winners’ are winners, and also provided insight into the upper limits of human physiological performance and the capacity of physiological systems to acclimatize [22]. Beyond human athletes, the physiological limits, capacities and acclimatization potential of other animals are beginning to unfold (*e.g.,* [1] [23–25] [26] [27]). Ultimately, measurement systems that can reliably and precisely estimate the repeatable physiological variation among individuals remain at the heart of both biomedical research and evolutionary biology [17], despite the attraction of “*The Golden Mean*” and a long tradition in scientific culture of minimizing individual variation in our data to obtain a clear picture.

A core challenge for experimental biologists examining the repeatability of individual traits is experimental noise, which exists even when laboratory experiments are highly controlled (Fig. 1). Indeed, if experimental noise is too great when a physiologically meaningful trait is measured and then remeasured for an individual, a Type II statistical error (false negative) can emerge in both longitudinal and cross-sectional studies (because experimental noise equally influences control and treatment groups). Thus, both longitudinal and cross-sectional physiological studies often assume that a trait in an individual is temporally stable over the timescale of the experiment [28].

**Fig. 1.**
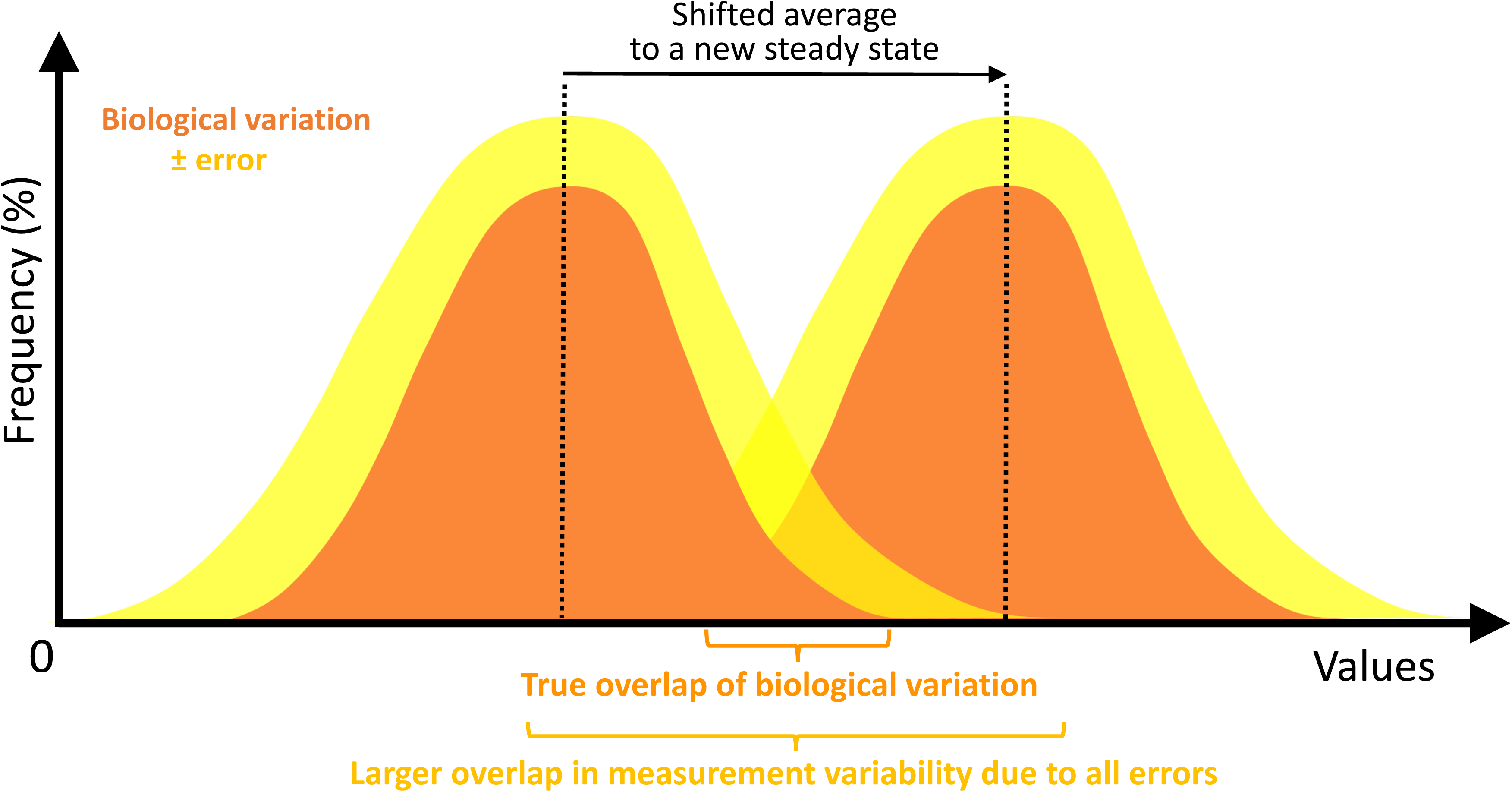
Schematic of the effects of methodological errors on measuring stable traits. When measuring a stable trait, the protocol, electronic, and analytical errors can inflate the variability of the measurement (for simplicity, the experimental noise is assumed to be equally distributed around the true mean in the schematic) and mask the real biological variation. At a new steady state, the distribution of biological variation can shift to a different level. If instrumental, protocol, and analytical errors are sufficient, measurement variability can overlap amply to compromise the precision in quantifying the real biological variation. The principal idea is to eliminate sources of error as best as possible to detect 1) what the real biological variation is and 2) any shifts in the true value of an individual metric.

Yet, physiological systems are dynamic, *e.g.*, ontogeny and aging [29]. Moreover, comparative physiologists who study ectotherms routinely measure this dynamic when measuring acclimation or acclimatization responses to new environmental conditions.

Therefore, while comparative physiologists have necessarily examined the temporal stability of a physiological trait in “control” animals, the understanding of the repeatability of physiological traits under “experimental” conditions (where the animal is challenged to perform) lags well behind that of sports and biomedical physiology [e.g., exercise performance [30] [31], temperature regulation [32] [33] [34] [35], and overall metabolism [36] [37] [38] [39].

Consequently, the major goal of this study is to propose and test an analytical approach that would help comparative physiologists better separate those traits within an individual that are repeatable, from those traits confounded by too much experimental noise. Herein, we propose a Precision-&-Repeatability Assessment Matrix (PRAM) that integrates established statistical methods that can measure variability and individual repeatability (*see* Glossary for important distinction). Furthermore, our case study, which focuses on whole-organism physiological performance, is a suitable candidate for all animals because of its established relationship to locomotion performance [40], its important implications on lifetime fitness [41] [42], and its demonstrated genetic underpinnings in some ectothermic vertebrate systems [27] [43].

In particular, metabolic rate, typically estimated by measuring oxygen (O_2_) uptake under various conditions (*e.g.,* basal, routine, or maximum sustainable O_2_ uptake rates) and environments, has been broadly used to assess the robustness of the vertebrate cardiorespiratory system [44] [45] and even linked to human longevity [46] [47] [48]. We also consider metrics of whole-animal anaerobic metabolism, the glycolytic pathways that utilize substrate-level phosphorylation to generate ATP, which in contrast, play a crucial role in shorter-term, life-promoting situations [6] [2] [49] [50] [51] [52] [53], such as environmental hypoxia [54] [2] [55], predation and predator-escape responses [56] [57] [58] [59], and difficult reproductive migration passages [60] [23]. Indeed, anaerobic metabolic metrics have received scant attention; just 12% of published repeatability studies in sports medicine and comparative physiology (summarized in Table S1) have examined indirect metrics to make largely inferential assessments of whole-animal anaerobic performance (Table S1). Here, we take advantage of the standardized Integrated Respiratory Assessment Protocol (IRAP, see supplementary materials for details) that measures both aerobic and anaerobic metabolic metrics on individual fish. No single study to date has compared the repeatability of aerobic and non-aerobic metabolic metrics. Therefore, our secondary goal is to compare the repeatability of aerobic and non-aerobic metrics for whole-animal metabolism measured by IRAP.

## The focus of comparative physiology on the metabolic rate of fishes

Comparative physiologists study perhaps the most diverse biological system among all experimental biologists, which in turn is well poised to tackle the challenges of studying biological variations. Among all the physiological performance traits studied in vertebrates, a great deal is known about the precision and reliability of whole-animal metabolic measurement techniques in fish [1] [6] [61] [62] [63] [4] [64] [65] [66] [67] [68] [10] [69] [70] [71] [72]. Moreover, fish species are the most diverse vertebrate lineage (over 30,000 species), representing over 50% of vertebrate species by number and inhabiting almost every aquatic environment on the planet [73] [74]. Their physiological systems, as a result, are replete with biological variation and acclimation potential. Even a fundamental physiological trait such as blood O_2_ transport can vary enormously among fish species [75] [76], which have the only known vertebrate lineage without haemoglobin (*e.g.*, icefish [77]). Within an individual, different haemoglobin isoforms can change over periods of three weeks [78], and factors that alter the oxygen affinity of hemoglobin can be modified within hours [79] [80].

Yet, comparative physiologists have continued to package individual variation around a mean value bounded by the standard error bars rather than exploring the potential significance of this individual variation, namely the ‘Tyranny of the ‘Golden Mean’ [17].

Although the reasons for the emphasis on the ‘Golden Mean’ in comparative physiology remain valid and multifaceted [81], gains are being made in acknowledging individual variation, reporting the precision of measurements [11] and reporting experimental conditions to better assess the methodologies [81]. (Table 1 summarizes the challenges of pursuing high-quality experimental studies of physiological performance alongside immediate and long-term solutions.). Therefore, we propose that comparative physiology and fish physiology specifically can benefit from the application of PRAM, which incorporates some of the key operational principles used in biomedical physiology and animal ecology.

**Table 1.**
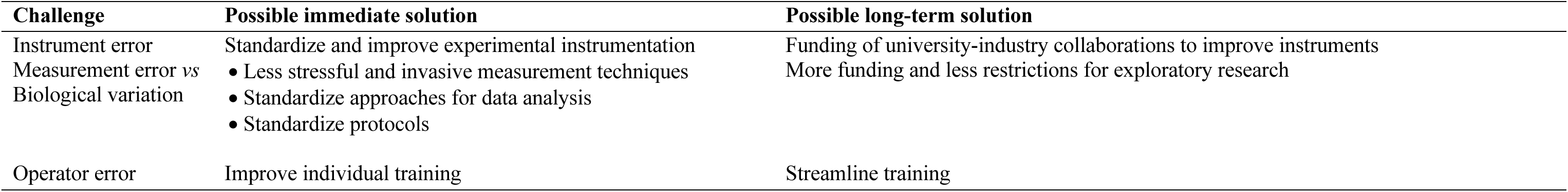
Challenges and solutions to improve data quality. Three general areas of challenge that collectively contribute to experimental noise are instruments, biology and operators. For each area, suggestions for both short- and long-term solutions are offered.

## Approaches used by other disciplines to account for individual variation

Biomedical physiologists typically identify a healthy human state within a range of values for a given parameter that is normal for individuals (rather than a single mean of a population), an approach not systematically explored by comparative physiologists [82] [11]. The presumption with this approach is that the parameter can be measured precisely.

A Bland-Altman analysis, a statistical approach to assess individual variation [3] [83], is often used in sports physiology. It calculates the percentage difference between two measurements taken from the same individual, divided by their average [3]. The percentage difference measured on the same individual and the average value become a single data point on a Bland-Altman plot (Fig. 2). Experimental noise, however, is incorporated into a Bland-Altman analysis of individual variation.

**Fig. 2.**
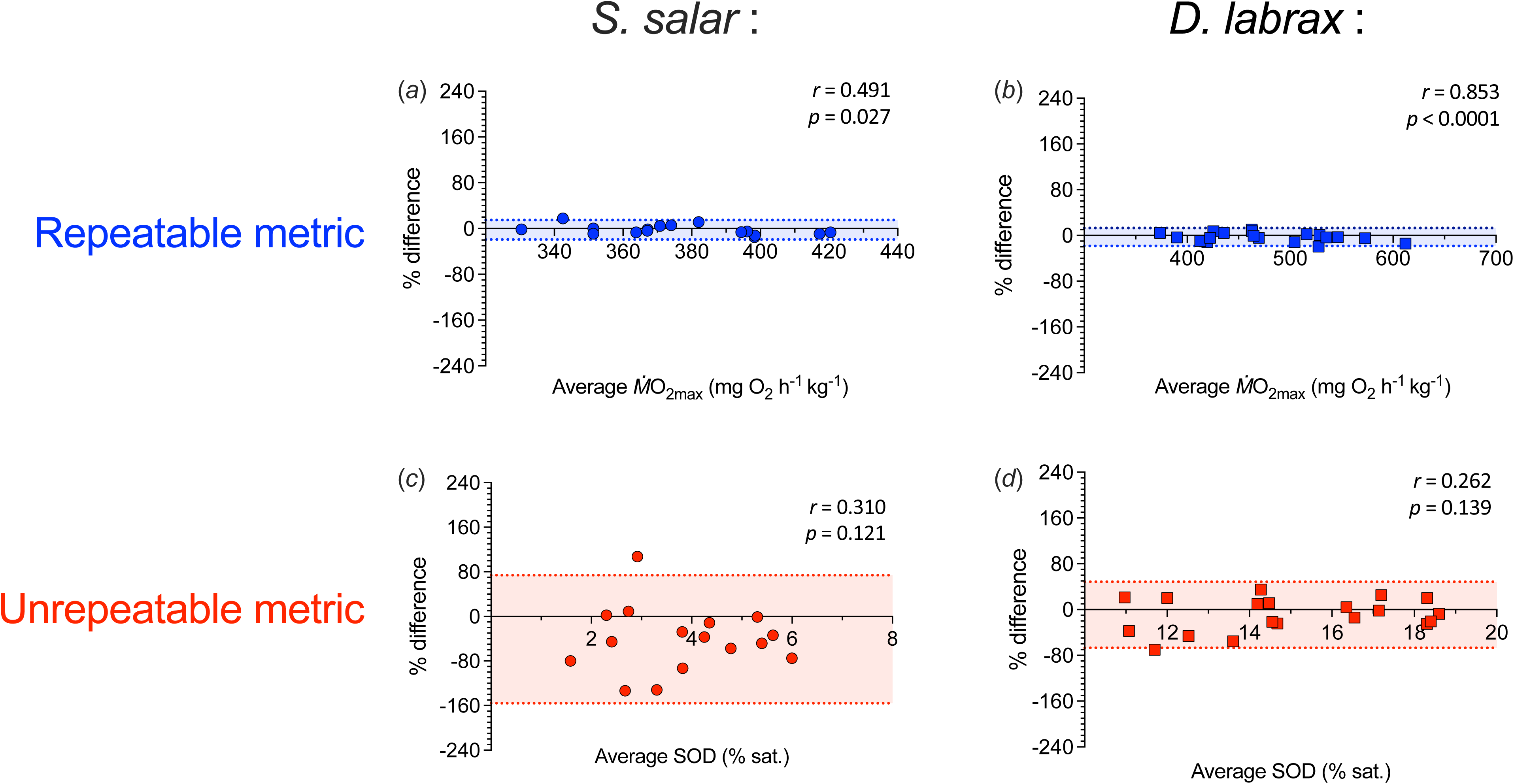
Bland-Altman analyses of repeatable and unrepeatable metabolic metrics characterized by the standardized Integrated Respiratory Assessment Protocol (IRAP), which measures whole-animal aerobic and non-aerobic metabolic metrics. Metrics were remeasured 4 weeks apart on the same individuals of European sea bass (*Dicentrarchus labrax*) at 25 °C (∼30 ppt, ∼60 g, n=16) and Atlantic salmon (*Salmo salar*) at 11 °C (∼30 ppt, ∼75 g, n=16). Metrics in blue were repeatable; metrics in red were unrepeatable (as determined by the Pearson correlation coefficient test using *p* < 0.05 (*r*-value: *p*-values appeared in the upper right corner of each figure). Coloured shading indicates 95% limits of agreement, a measure of the metric variability (twice its standard deviation and accounting for the mean difference of the two measurements). Here, as representatives, an aerobic metabolic metric is the maximum oxygen uptake rate (*Ṁ*O_2max_) and a non-aerobic metabolic metric is scope for oxygen deficit (SOD). The repeatable aerobic metabolic metric has a much narrower band of intra-individual variability (a smaller value of 95% limits of agreement) than the unrepeatable non-aerobic metabolic metric. The data were adapted from the published studies [98] [100].

The normalized coefficient of repeatability (COR) for a group of individuals quantifies the variability of a metric (Fig. 2) (Eqn. 1) [83].

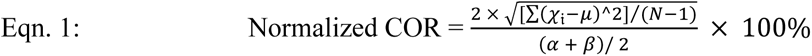

where *χ_i_* is each value of the difference between two repeated measurements from each individual, *μ* is the mean of the difference between the two repeated measurements, N is the sample size, *α* is the value of a metric measured in the 1^st^ test, and *β* is the value of a metric measured in the 2^nd^ test.

Repeatability of rank order in a repeated measurement design has been used by comparative physiologists to examine if the individuals maintain their relative rank within a population despite a shift in the population average (Also see the rank order analyses of our dataset in Figs. S1, S2) [84] [85] [86] [87] [88] [89]). Analysis of rank order, however, loses the quantitative aspect of the metric under consideration: the winner can remain the winner, but information on the absolute performance is lost.

Generalized Linear Mixed Models (GLMM) are often used by animal ecologists and ecophysiologists working in field settings who face a greater challenge than laboratory-based experimental biologists concerning the ability/inability to control important environmental variables, e.g., temperature, rainfall, humidity, salinity, oxygen levels, and photoperiod.

Uncontrolled environmental variables add to experimental measurement “noise”. In this case, a Generalized Linear Mixed Model (GLMM) statistical approach can account for the impacts of variables that cannot be controlled [90] [91] [92] and the contribution of the experimental noise [93] [94] when testing the repeatability of measured metrics. A GLMM, however, does not attempt to separate experimental measurement noise from true individual variation.

## Precision-&-Repeatability Assessment Matrix (PRAM)

In view of the limitations for each of the statistical approaches noted above (Table S2), we propose a Precision-&-Repeatability Assessment Matrix (PRAM) that graphically combines a parametric test of repeatability (the Pearson correlation coefficient) with a normalized COR. This theoretical framework then provides a quantitative approach for experimental biologists to better visualize the repeatability and variability of metrics (Fig. 3).

**Fig. 3.**
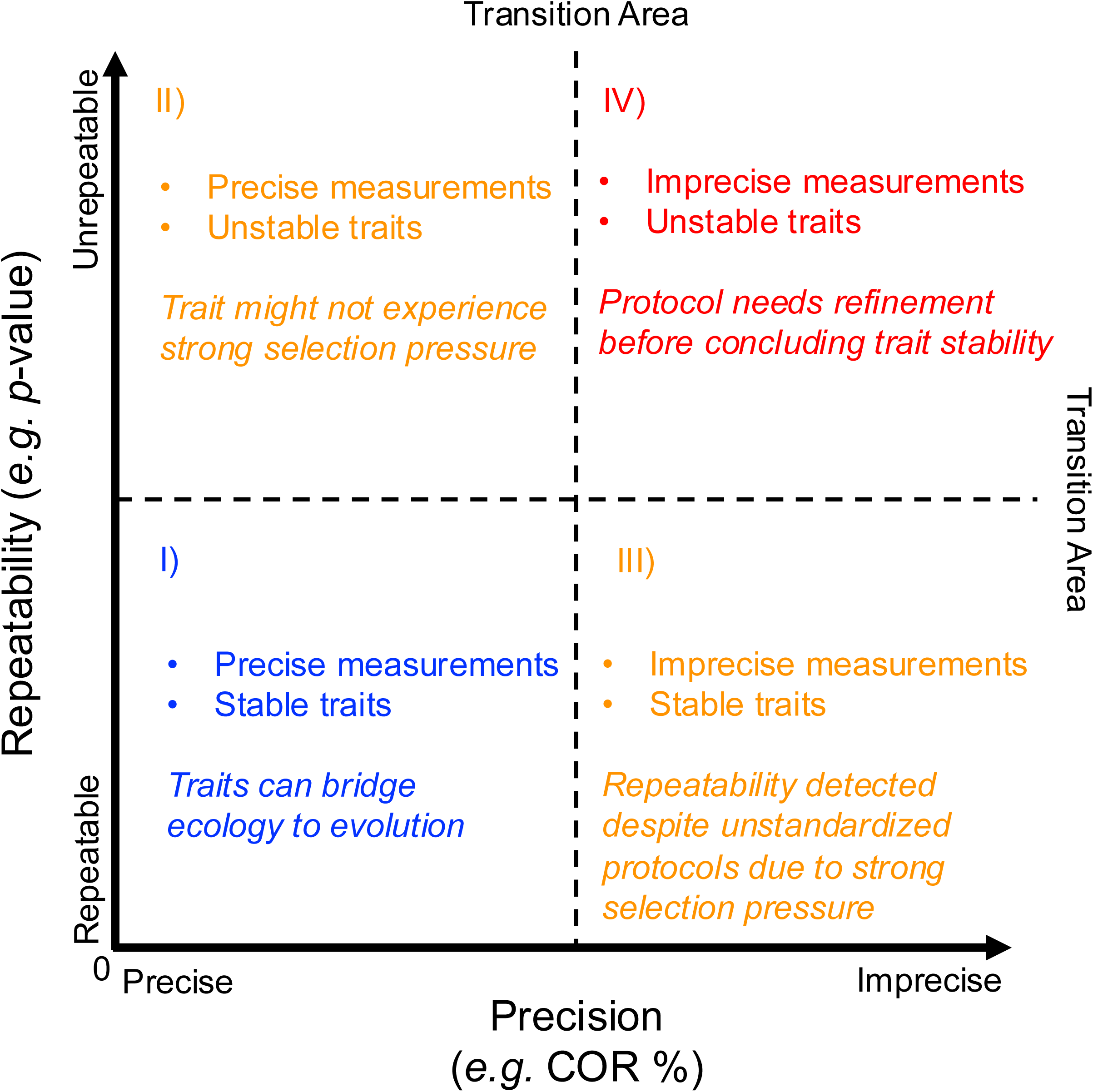
Theoretical framework for measurement precision and trait stability using a Precision-&-Repeatability Assessment Matrix (PRAM). The decision matrix is an analytical concept and approach based on the association between the variability of metrics and the repeatability of the metrics. The utility of the decision matrix is to distinguish the stability of traits in individuals from the experimental noise. The normalized coefficient of repeatability (COR) is a proxy for measurement precision at the x-axis, and the *p*-value from correlation coefficient tests is a proxy for repeatability at the y-axis. This forms quadrants of interest: i) precise measurements and stable traits; ii) precise measurements and unstable traits; iii) imprecise measurements and stable traits; iv) imprecise measurements and unstable traits. The area of transition for repeatability is a *p*-value of 0.05 using correlation coefficient tests. The area of transition for precision is the normalized coefficient of repeatability value with threshold values corresponding to a higher probability for the metrics to become repeatable (*e.g. p*-value < 0.05).

PRAM statistically quantifies variability within the individual-based measurements using the well-established Bland-Altman analysis [95] [3] [96] [30]. Also, correlation analysis (*e.g.* Pearson’s correlation coefficients) assesses the repeatability of a given metric measured twice on the same individual using a balanced design [97] [90]. These two statistical analyses are then placed on Cartesian coordinates to arrive at a decision matrix with four quadrants (Fig. 3) that inform the quality of the performance metric by parsing out the biological variation among a group of individuals and experimental noise. For example, the quadrant closest to the origin identifies a metric that is stable and measured by precise protocols (*i.e.,* resulting in low experimental noise). These repeatable metrics would be best suited for testing hypotheses, developing theories, and building models with strong predictive power. Moving away from the origin of the quadrant, the other quadrants represent metrics that are less repeatable for different reasons, *i.e.,* precise measurements applied to unstable traits, imprecise measurements applied to stable traits, and imprecise measurements applied to unstable traits (Fig. 3). Therefore, the decision matrix of PRAM helps distinguish experimental noise from individual variation, enabling a better understanding of physiological variation.

## A case study: applying PRAM to whole-animal metabolic metrics for fish

The data used for the case study were those collected from two independent repeated-measure design studies with two genetically distant athletic fish species, Atlantic salmon (*Salmo salar*) and European sea bass (*Dicentrarchus labrax*) [98] [99] [100]. Both studies retested the same individuals after ∼4 weeks (*see* Supplementary Materials for animal information). Moreover, both used IRAP, which holistically characterizes an individual fish’s respiratory phenotype by measuring up to 13 metrics for the performance and capacity of aerobic and anaerobic metabolism [8] (*see* Glossary for full definitions of terms & supplementary material for details of their measurement, Fig. S3, S4). Consequently, the same practitioners performed the standardized IRAP procedure under controlled laboratory conditions and with similar equipment (Table S3), which minimized operator variability (*see* Table 1). Also, phenotype stability assessment was conducted over a 1-2 month time frame, acknowledging that fish can partially or fully acclimate to new environmental conditions over several weeks [101] [102] [103] and that only similar life stages should be compared in the first instance.

In principle, the temporally stable traits measured precisely should be repeatable across species. Otherwise, the targeted trait either does not have a consistent stability among individuals within in species, or the measurement is imprecise. PRAM was then applied to identify which individual traits were stable, which were not and which were noisy (Fig. 4).

**Fig. 4.**
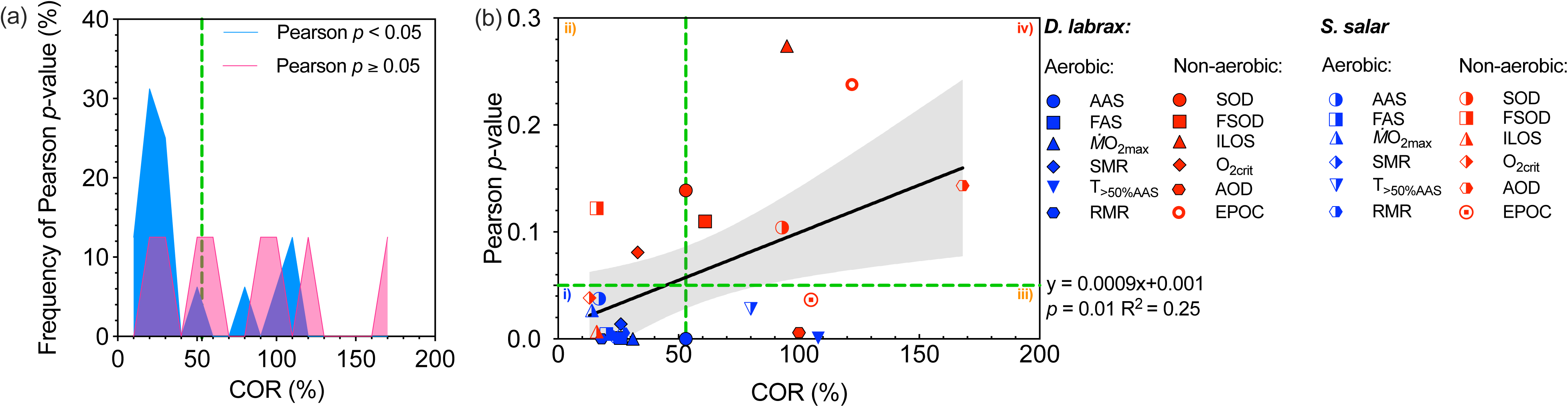
An examination of the relationship between the variabilities of metabolic metrics with the repeatability of the metabolic metrics. (a) The frequency distributions of the *p*- value of Pearson correlation coefficient tests (categorized into groups of Pearson *p* < 0.05 and Pearson *p* ≥ 0.05) informed the formation of quadrants in the Precision-&-Repeatability Assessment Matrix (PRAM; *see* Fig. 3). The green vertical dashed lines [normalized coefficient of repeatability (COR) = 53%] are based on the frequency distribution of the *p*- value of Pearson correlation coefficient tests. When the metrics have COR values of less than 53% (between the origin and the green vertical dashed lines), the metrics have nearly three times higher probabilities of being repeatable (*p* < 0.05). When COR is larger than 53% (beyond the green vertical dashed lines), the metrics had a lower chance of being repeatable. (b) The green vertical dashed lines annotate the threshold (COR = 53%) where metrics change from being precise to imprecise. The *p*-value of 0.05 is annotated by the green horizontal dashed lines to indicate the area where the metrics gradually become unrepeatable from being repeatable. Using these arbitrary threshold values of repeatability and precision, quadrants were formed in panels using the theoretical framework of PRAM: i) precise measurements and stable traits; ii) precise measurements and unstable traits; iii) imprecise measurements and stable traits; iv) imprecise measurements and unstable traits. Different symbols denote different metabolic metrics. To understand the general relationship between variabilities and repeatability, the analyses combined the measurements obtained from European sea bass (*Dicentrarchus labrax*) at 25 °C (∼30 ppt, ∼60g, n=16; filled symbols) and Atlantic salmon (*Salmo salar*) at 11 °C (∼30 ppt, ∼75g, n=16; half-filled symbols). Metrics in blue measure traits when animals predominantly rely on aerobic metabolism (*i.e.* aerobic traits), and metrics in red measure traits when animals engage in a substantial amount of glycolysis (*i.e.* non-aerobic traits). Aerobic metabolic metrics (in blue) include absolute aerobic scope (AAS), factorial aerobic scope (FAS), maximum oxygen uptake (*Ṁ*O_2max_), standard metabolic rate (SMR), routine metabolic rate (RMR), and time spent above 50% AAS (T_> 50% AAS_). Non-aerobic metabolic metrics (in red) include excess post-exercise oxygen consumption (EPOC), scope for oxygen deficit (SOD), the factorial scope for oxygen deficit (FSOD), incipient lethal oxygen saturation (ILOS), accumulated oxygen deficit (AOD) and critical oxygen saturation (O_2crit_). In general, less variability in measuring the metrics is positively related to the higher probabilities of being repeatable metrics. The data were adapted from the published studies [98] [100].

When COR was less than 53%, a metric was nearly three times likely to be repeatable (the peak of the frequency distribution of arbitrary *p*-value < 0.05 was nearly triple the peak of the frequency distribution of *p*-value ≥ 0.05, Fig. 4a). When COR was greater than 53%, the chance of a metric being unrepeatable increased (Fig. 4a). Therefore, we used COR of 53% as a nominal line of reference for the measurement precision in this analysis (Note: 53% is applicable for this dataset and a work in progress). The working criteria to set the threshold value are 1) the transition point in the frequency distribution of COR where metrics start to shift from a higher chance (e.g. 2-3 times more likely) being repeatable to a lower (e.g. 0.5 times) chance being repeatable. We then set the quadrants for PRAM arbitrarily as COR of 53% for measurement precision (green vertical dashed line on the x-axis) and repeatability (green horizontal dashed line on the y-axis) (Fig. 4b). As perhaps expected, the more precisely measured metrics (*i.e.* smaller COR values) tended to be more repeatable (correlation test: *p*-value ≤ 0.01, Fig. 4b), but with some exceptions, *e.g.,* FSOD was measured precisely and was not repeatable (Fig. 4b).

PRAM also demonstrated that individual aerobic metabolism traits (AAS, FAS, *Ṁ*O_2max_, SMR, and RMR) were repeatable for both species of athletic fish. The only unrepeatable aerobic metric was T_>50%AAS_, an index of spontaneous activity that is likely context-dependent. *Ṁ*O_2max_ is an important independent metric to derive AAS & FAS. Bland-Altman analysis on *Ṁ*O_2max_ demonstrated that *Ṁ*O_2max_ has a limited measurement variability (the narrow horizontal band in Fig. 2; Bland-Altman analyses of other metrics are in Fig. S5 & S6), a finding that may reflect past efforts to establish reliable measurement techniques for *Ṁ*O_2max_ [10] [69] [104] [45] and the tightly coupled steps of the O_2_ transport cascade for peak performance in vertebrates [105] [106]. When measured properly, *Ṁ*O_2max_ should have limited intra-individual variation in a repeated measure design, as previously shown in healthy humans [83] [107] [106] [22]. Thus, PRAM analysis confirmed that measurement precision and trait stability were necessary features of the well-established independent performance metrics. Indeed, studies with other animal taxa find strong repeatability of aerobic metabolic metrics (Table S1). Also, SMR, *Ṁ*O_2max_, AAS and FAS have been related to the lifetime fitness of vertebrate species in various environments [108] [109] [110] [111] [27], with some but not all [112] studies finding stable traits within a life stage.

A novel discovery of the PRAM analysis was that whole-organism anaerobic metabolism metrics were generally either unstable or were imprecisely measured. For example, EPOC, a metric that predominantly measures the oxygen equivalents of glycolytic metabolism during exhaustive exercise, was reported as both a stable and an unstable metric. This discovery may reflect the many challenges that still exist to properly quantify anaerobic metabolism at the organismal level (*see* reviews [113] [114] [115] [116]), those beyond the extent to which glycolysis is affected by individual variation in cardiorespiratory robustness [71] [72] [2] [117] [100]. For example, differences in phosphagen and metabolic substrate stores, as well as glycogenesis, can all potentially introduce variation [118] [119] [120]. Similar concerns also apply to the scope for oxygen deficit, a new performance proxy for the anaerobic capacity of a fish in a hypoxia challenge test, [8] which quantifies the accumulation of glycolytic end-products and different demands on glycolysis in severe hypoxia [8] [121].

The Bland-Altman analysis for the scope for oxygen deficit revealed large variability (a wider horizontal band in Fig. 2; see Bland-Altman analyses for other metrics in Fig. S5 & S6). Thus, assessment protocols for whole-animal anaerobic capacity and performance, which remain in their infancy, perhaps should be given greater attention to develop more precise methods, especially given that they fuel shorter-term, life-saving activities.

Surprisingly, PRAM analysis revealed important nuances for two widely used and well-established metrics used to estimate hypoxia tolerance in fish, *i.e.,* O_2crit_ and ILOS, which were measured with the same standardized hypoxia challenge test [9]. While O_2crit_ was precisely measured (*i.e.* low experimental noise) in both species, ILOS was precise only in Atlantic salmon (*Salmo salar*) and not in European sea bass (*Dicentrarchus labrax*) (Fig. 4b). O_2crit_, as calculated here, benefits from an objective mathematical approach (SMR is the baseline to derive an intercept with a linear regression equation [64] [5]). In contrast, ILOS is a visually subjective recording of the ambient O_2_ level when a fish loses its upright equilibrium. Why O_2crit_ demonstrated inter-specific variation [67] for its repeatability between *S. salar* but not *D. labrax* is unclear.

## Future directions for research on the repeatability of individual variation in experimental biology

Our case study of PRAM using IRAP metrics for two species of ray-finned fish is only a demonstration that informs us of a few key general principles for experimental biology. An unstable trait can be unrepeatable even when precisely measured (Quadrant II in PRAM; Fig. 3). Conversely, and as expected, a stable trait requires a standardized protocol to generate precise measurements (*i.e.,* low experimental noise) (Quadrant III in PRAM; Fig. 3). Being able to distinguish the two scenarios is important for biological experiments, and this is what PRAM achieved. The same principles likely apply to other performance metrics in other animal and human models, and perhaps across multiple biological organizations.

Striving for repeatability has been a common goal among experimental biologists in recent literature [122] [123] [124] [125] [11]. The central idea behind the PRAM is to characterize the repeatability and measurement variability of a metric on a decision matrix to distinguish the true physiological variation from experimental noise. PRAM assesses the quality of the metrics and informs the types of conclusions that can be drawn. Much work remains, however, to identify the multifaceted physiological reasons for the differences in repeatability for metabolic metrics, as well as further analyses of the PRAM framework in acclimation (or acclimatization) studies. For example, how certain acclimation states can lead to less stable traits (e.g. transition from quadrant I to quadrant II). Hence, we welcome future improvements to the decision matrix, given that statistical assessment methods are constantly evolving. For example, the quantification of variance can use the coefficient of variation or covariance [126] [91], and the assessment of repeatability can use residual variance [90]. For repeatability tests conducted on more than two time points (*e.g.* [127] [58] also see reviews on this topic: [126] [128]), the decision matrix can identify the sources of the variability in a pair-wise manner across multiple time sampling points. The criteria for the COR threshold that determine a precise measurement are a work in progress. It remains to be tested with future studies whether a more rigid COR threshold can be established (*e.g.*, ∼50% COR).

The PRAM framework can also contribute to two long-standing questions in experimental biology: 1) How to mitigate the trade-offs between sample size and measurement precision to improve statistical power at a limited experimental resource? 2) How to account for the learning behaviours over multiple repeatability tests? First, if the imprecise measurement (quadrant III) quantifies the stable traits, a large sample size can potentially compensate for the statistical power. Still, increasing the sample size of imprecise measurement would be less productive than trying to source and reduce the noise. If the measurements are precise and targeting stable traits (quadrant I), a smaller sample size might be needed to reach the desired statistical power. The precise metrics would better utilize the limited research resources when quantifying the plastic traits (*e.g.*, quadrant II). Moreover, our PRAM framework could be used to retrospectively analyze existing data, as shown here, to identify metrics that appear robust (quadrant I) and in need of some scrutiny (quadrants III & IV), as well as either rejecting those metrics in quadrant IV or investigating the underlying issues. One potential issue worth investigating is the importance of learning behaviour in the quality of a metric. Using the PRAM framework as part of an experimental design can help answer the question of how many repeatability tests and over what time intervals are necessary.

Overall, PRAM analysis will help comparative physiologists and experimental biologists to objectively, quantitatively, visually and more systematically understand the metrics they measure. PRAM can be applied well beyond the limited number of metrics used here as a case study. Furthermore, PRAM can better calibrate the confidence placed in the indicators that we believe are metrics of fundamental metabolic traits. This analytical approach can enable physiological metrics to be better integrated into studies of genomics, molecular mechanisms, biomechanics, the life history of the organisms, and the environmental selection pressure over the evolutionary history. After all, inter-individual variation is the raw material for evolution through natural selection.

## Ethics

This work has ethical approvals from animal welfare committees (*see* supplementary methods for the certificate numbers of the animal welfare approvals).

## Data accessibility

Data are archived in a repository for publication.

## Acknowledgments

Many thanks to members of the Department of Zoology at the University of British Columbia and George Lauder at Harvard University for numerous discussions about individual variation and the repeatability of metrics. To Dave Randall’s question of what defines a trait.

## Author contributions

Y.Z., C.M.W., C.J.B., A.P.F. conceptualized the study. Y.Z. performed experiments and data analyses, and wrote the original manuscript. All authors provided manuscript edits and comments and approved the final version.

